# Geometric averaging provides normalization-invariant feature ranking in compositional sequencing data

**DOI:** 10.64898/2026.05.16.725171

**Authors:** Emilia Nunzi, Luigina Romani

## Abstract

In compositional next-generation sequencing (NGS) analyses (including microbiome studies, RNA-seq and metagenomics) the arithmetic mean (AM) of relative proportions is the default operator for summarizing feature abundances. We show that this default produces unstable rankings in real compositional data. Across 102 prevalent genera in the dietswap dataset (n=38 baseline samples), 23 genera (22.5%), including members of *Bacteroides*, *Eubacterium* and *Bilophila*, yielded opposite group-level conclusions under AM and the geometric mean (GM).

This pattern reflects two formal properties of compositional aggregation. First, AM-based rankings change with the within-sample normalization domain, whereas GM-based rankings are invariant under the multiplicative structure of compositional data. Second, the centered log-ratio (CLR) transformation absorbs geometric averaging into the data representation, so that arithmetic averaging on CLR-space recovers the GM ranking exactly. Both properties were verified numerically on the dietswap dataset, where the Spearman correlation between GM- and CLR-based rankings was 1.000 in both groups.

The operator-choice problem propagates to between-group differential inference: under AM, log_2_ fold-changes vary across normalizations and the relative ranking of features by effect size is not preserved; under GM and CLR, the ranking is preserved. We recommend GM-based summaries for feature ranking and CLR-transformed abundances for cross-sample comparisons. This change requires no new computational tools and is fully compatible with existing differential-abundance pipelines, but eliminates an under-recognized source of irreproducibility in biomarker discovery across microbiome studies, transcriptomics, metagenomics, and mass-spectrometry-based metabolomics, in all settings where features are quantified relative to a sample total.

**IMPORTANCE:** Studies of the gut microbiome routinely identify which bacterial groups are more or less abundant in patients versus healthy controls, in different diets, or before and after a treatment. The same kind of comparison underlies sequencing-based analyses across biology, from gene expression to metagenomics. To do this, researchers must average the abundance of each measured entity across many samples, and the standard choice is the simple arithmetic average. We show that this choice can be misleading for any data where each measurement is expressed relative to a sample total, as is typical of sequencing-based assays, and that in real data it can flip the answer to which group is more enriched. Analyzing a published dietary intervention study, we found that one in five gut bacteria (including *Bacteroides* and *Eubacterium*) gave opposite results depending on which average was used. Switching to the geometric average resolves this inconsistency and makes biomarker discovery more reproducible. This change is immediate to implement (it does not require new software or specialized training) and applies not only to microbiome studies, but to any biological measurement where what is detected, whether a gene transcript, a microbial taxon, or a metabolite, is quantified relative to a sample total: gene-expression analysis, metagenomics, and metabolomics among others.

## INTRODUCTION

Across biology, quantitative profiling technologies share a fundamental constraint: the count or intensity of each feature (a microbial taxon, a gene transcript, a metabolic pathway, or a metabolite), whether produced by 16S rRNA amplicon sequencing, RNA-seq, shotgun metagenomics, single-cell transcriptomics, or mass-spectrometry-based metabolomics, is meaningful only relative to the sample total, since that total is fixed by sequencing depth or extraction yield rather than by the underlying biological load. Data with this property are called compositional [6, 2, 21]. Within-sample (WS) normalization is therefore necessary to render abundances comparable, and the choice of normalization (to total reads, to a reference feature, to a spike-in standard) influences which features appear enriched and which depleted [22, 3, 5, 11, 12].

Beyond WS normalization, a second choice shapes the analysis: how to summarize feature abundances across the samples of a group, for example when identifying which microbial taxa are most enriched in a diseased cohort, which gene transcripts characterize a treatment condition, or which metabolites distinguish two dietary groups. Across-sample (AS) summarization typically defaults to the AM of relative abundance or ratios. The AM, however, is sensitive to the total number of counts in each sample: high-depth samples carry more weight in the group average, even for features whose relative abundance is the same across all samples, thus introducing instability in the feature ranking. The GM, by contrast, operates in log space, preserves multiplicative relationships, and is invariant to within-sample rescaling [15]: a feature’s GM-based group summary depends only on its relative abundance in each sample, and is unaffected by differences in sequencing depth across samples.

The choice of averaging operator at this aggregation step is logically prior to differential-abundance testing: any procedure that relies on AM-based summaries of raw proportions or counts inherits their normalization dependence. Despite this, AM-based summaries remain the default in much of the applied microbiome literature, and the practical consequences of this default for biomarker discovery have not been characterized on real data. A natural solution is to work in log space, where arithmetic averaging recovers the geometric mean on the original scale. This is precisely what the CLR transformation [6] achieves: by expressing each feature relative to the WS-GM before averaging, CLR-based summaries are normalization-invariant by construction. Established differential-abundance methods such as ANCOM-BC [10] and LinDA [23] implicitly adopt this approach, operating on CLR-transformed abundances rather than raw proportions.

In this work, we (i) formalize the conditions under which AM- and GM-based aggregation produce different feature rankings on compositional data (Text S1); (ii) illustrate the resulting normalization dependence on minimal worked examples; and (iii) quantify the empirical extent of AM/GM discordance on a publicly available dietary intervention dataset (dietswap, 102 prevalent gut microbial genera [24]), showing that 22.5% of these genera yield opposite group-level conclusions under AM and GM, and that GM-based rankings coincide exactly with arithmetic averaging on CLR-transformed data. While our empirical demonstration uses 16S microbiome data, the framework applies to any compositional NGS measurement and, more broadly, to any assay producing relative abundance vectors, including metabolomics. Throughout, our focus is on feature ranking under compositional constraints, not on differential abundance inference, and we treat zero handling as a separate concern that can be addressed by sparsity-aware methods downstream.

## RESULTS

### Geometric mean provides normalization-invariant feature ranking in compositional data

In comparative microbiome and genomic analyses, identifying which features are consistently more or less abundant across sample groups is a prerequisite for defining microbial or genetic signatures. Because NGS data are compositional, feature abundances acquire biological meaning only after normalization to a sample-specific reference (total reads, a target feature, the WS-GM, …), and the choice of reference directly shapes downstream comparisons.Here we examine how the choice of summary operator (AM vs. GM) on sample groups interacts with the choice of per-sample normalization for the purpose of AS feature classification. Two formal properties underlie the analyses that follow (Text S1): (i) AM-based rankings change with the WS-normalization domain, whereas GM-based rankings are invariant by the multiplicative property GM(*a*/*b*) = GM(*a*)/GM(*b*); (ii) CLR transformation absorbs geometric averaging into the data representation, so that arithmetic averaging across samples on CLR space recovers the GM ranking exactly. We illustrate both properties on minimal worked examples (Figure **1**–Figure **2**) before validating them on real microbiome data in the next sections.

**FIG 1.**
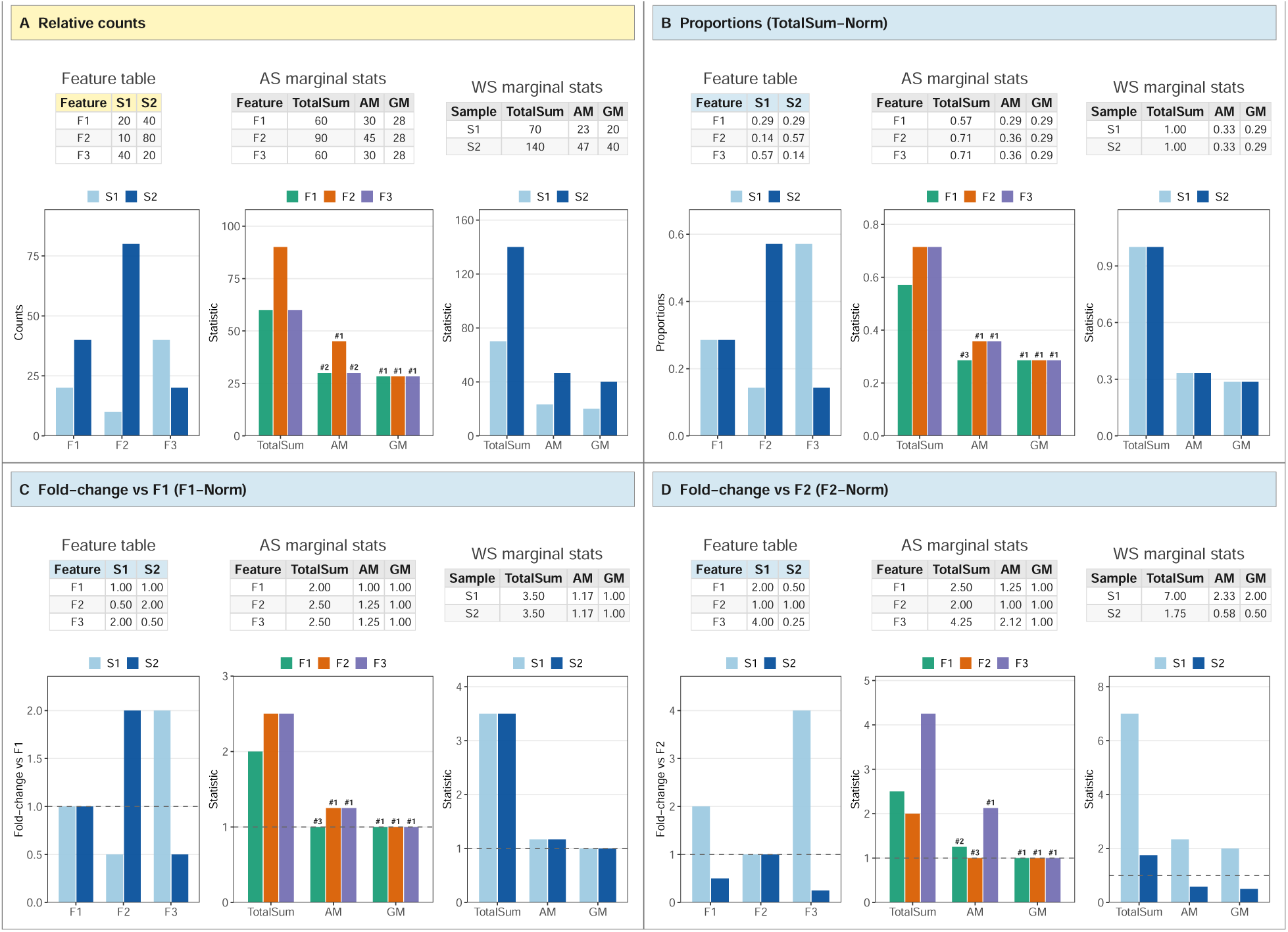
Two-sample illustration of normalization-dependent feature ranking under arithmetic versus geometric mean aggregation. Three features (F1–F3) measured in two samples (S1, S2). Each panel shows the feature table (yellow = raw counts in panel A; light blue = within-sample (WS) normalized values in panels B–D) with the corresponding across-sample (AS) and within-sample marginal statistics (light gray): total sum (Sum), arithmetic mean (AM), and geometric mean (GM). Bar charts plot the values from the corresponding table above. Panel A: relative counts. Panel B: proportions (counts normalized to the WS total). Panel C: ratios to F1 (fold-changes relative to F1). Panel D: ratios to F2 (fold-changes relative to F2). Rank labels (#1, #2, #3) above the AM and GM bars in the AS-marginal-stats charts indicate the feature ordering induced by each operator; AS-AM rankings change with the normalization (compare #1, #2, #3 across panels) whereas AS-GM rankings remain identical (all three features tied at #1 across panels B–D). Formal statement and proof sketch are provided in Text S1.

**FIG 2.**
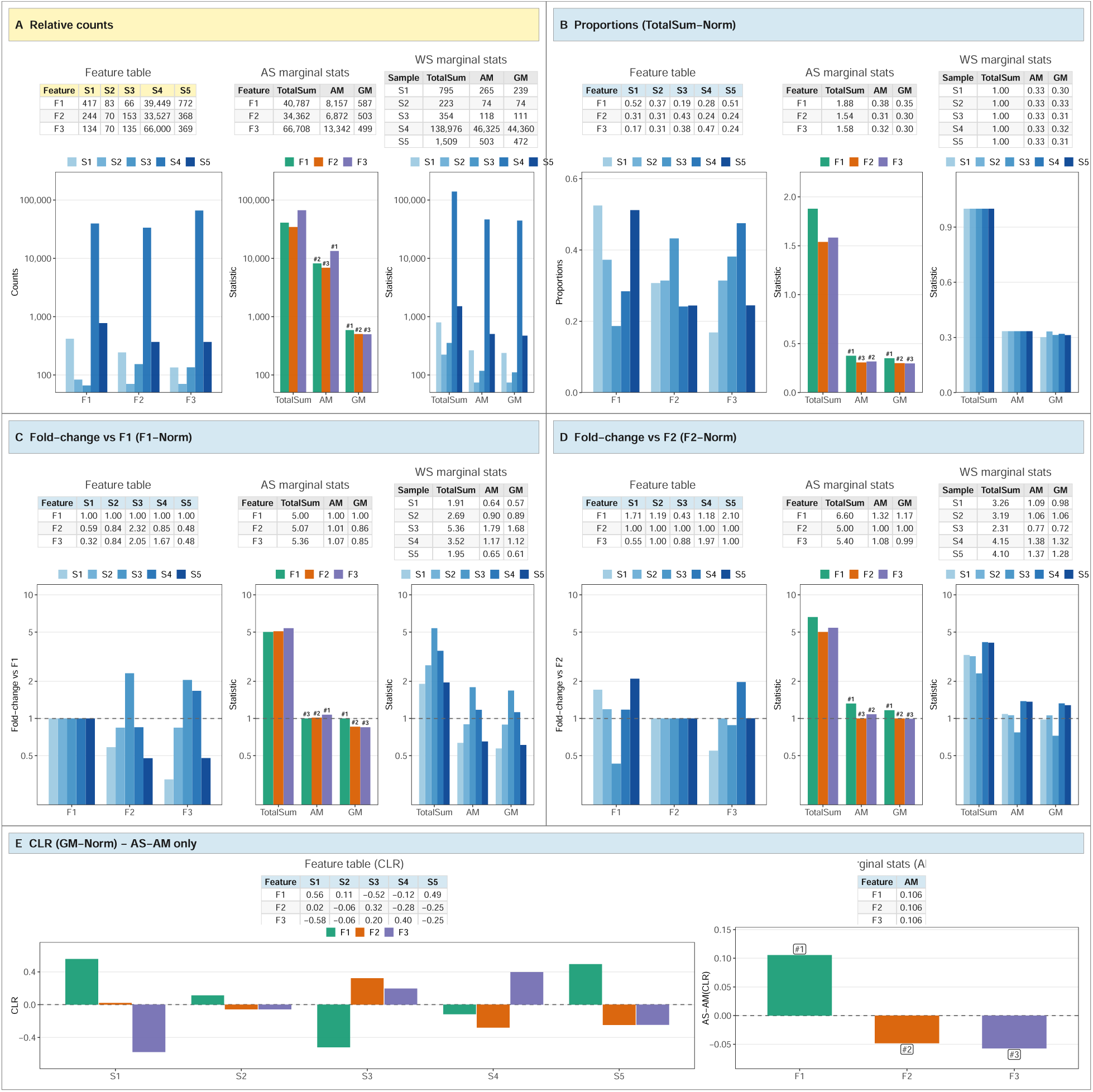
Robustness of geometric-mean-based ranking under highly uneven sequencing depths, including CLR transformation. Five samples (S1–S5) with sequencing depths spanning three orders of magnitude (S2 = 223 reads to S4 = 138,976 reads) and three features (F1–F3). Panels A–D each show the feature table (yellow = raw counts in panel A; light blue = WS-normalized values in panels B–D) with the corresponding marginal (AS, WS) statistics (light gray): total sum, AM, and GM. Bar charts plot the values from the corresponding table above. Panel A: relative counts. Panel B: proportions (counts normalized to the WS total). Panel C: ratios to F1 (fold-changes relative to F1). Panel D: ratios to F2 (fold-changes relative to F2). Rank labels (#1, #2, #3) above the AM and GM bars in the AS-marginal-stats charts indicate the feature ordering induced by each operator. AS-GM yields a stable feature ranking F1 > F2 ≈ F3 across all four normalizations (rank labels remain identical across panels), whereas AS-AM-based rankings change with the choice of reference and are dominated by the high-depth sample S4. Pairwise ratio comparisons are detailed in Table S1. Panel E: centered log-ratio (CLR) transformation of the same dataset, CLR(*x_i_*) = log(*x_i_*/GM(**x**)). The left part shows the per-sample CLR feature table and the corresponding bar chart (each sample’s CLR values sum to zero by construction); the right part shows the across-sample arithmetic mean AS-AM(CLR) per feature with rank labels. The AS-AM(CLR) ranking F1 > F2 ≈ F3 reproduces the AS-GM ranking of panels B–D, illustrating Proposition 2 of Text S1: across-sample arithmetic means of CLR values are equivalent (up to an additive sample-wide constant) to logarithms of AS-GM(proportions).

### Two-sample illustration of normalization dependence

Figure **1** considers three features (F1–F3) measured in two samples (S1, S2) under four equivalent representations: raw counts (panel A), proportions (panel B), and ratios to F1 or F2 (panels C–D). The feature composition shifts markedly between samples: F3 dominates S1 (40 counts, 57%) while F2 dominates S2 (80 counts, 57%), with F1 intermediate in both (20–40 counts, ∼29%). No feature is consistently enriched across the two samples; rather, the differences reflect a redistribution of relative abundances driven by sample-specific compositional shifts.

Despite these large changes in relative composition, AS-GM values are essentially identical for the three features and remain stable across all four normalizations (AS-GM ≈ 28 on counts; ≈ 0.29 on proportions; 1.00 on F1-ratios; 1.00 on F2-ratios), reflecting the multiplicative property GM(*a*/*b*) = GM(*a*)/GM(*b*). In contrast, AS-AM yields different feature rankings under each normalization. In panel D (ratios to F2), the AM-based summaries imply that F1 is approximately 25% more abundant than F2 (AS-AM = 1.25) and that F3 is more than twice as abundant as F2 (AS-AM = 2.12), even though the underlying counts of F2 (total = 90) exceed those of F1 (60) and F3 (60). The same caveat applies to the total sum, which is directly proportional to AM (TotalSum = *N*·AM) and therefore equally sensitive to depth-driven distortions. The GM-based summary, by contrast, returns 1.00 for all three features, correctly reflecting that no feature is consistently enriched relative to F2 across the two samples. The AM-based conclusion therefore depends on which feature is chosen as the reference, whereas the GM-based conclusion does not.

### Robustness to sequencing-depth heterogeneity

Figure **2** extends the same construction to five samples with strongly uneven sequencing depths (S2 = 223 reads to S4 = 138,976 reads), a regime more representative of real NGS datasets. The compositional structure is non-trivial: when expressed as proportions, F1 dominates in S1 (52.5%), F3 dominates in S4 (47.5%), while F2 shows moderate variation across samples (24–43%). On raw counts (panel A), AS-AM ranks F3 highest (13,341 reads), driven almost entirely by S4; AS-GM, by contrast, ranks F1 > F2 ≈ F3 (587, 503, 498), a ranking preserved when the data are expressed as proportions (panel B; AS-GM: F1 = 35.1%, F2 = 30.0%, F3 = 29.8%, versus AS-AM: F1 = 37.6%, F2 = 30.8%, F3 = 31.7%, with the F2/F3 ordering flipped under AM relative to GM), as F1-ratios (panel C), or as F2-ratios (panel D). The AM ranking instead changes with the normalization, and AM-based fold-change estimates are inconsistent under reference change: F2 appears 2% more abundant than F1 when normalizing to F1, but F1 appears 32% more abundant than F2 when normalizing to F2, a pair of estimates that cannot both be correct. The same caveat applies to the total sum, which is directly proportional to AM and therefore equally sensitive to depth-driven distortions. GM-based fold-changes are mutually consistent by construction (GM(F1/F2) = 1/GM(F2/F1); see Text S1 and Table S1 for the full set of pairwise comparisons). Together, panels A–D show that AS-GM provides a depth-agnostic estimate of typical relative abundance, while AS-AM is dominated by samples with extreme depth.

The multiplicative property of the GM provides a direct numerical verification: GM(F1/F2) computed on raw counts equals GM(F1)/GM(F2) = 586.79/503.13 = 1.17, identical to the GM computed directly on the F1/F2 ratios across samples (panel D). The same property gives the reciprocal identity 1.17 = 1/0.86, confirming that GM(F1/F2) = 1/GM(F2/F1) (Proposition 3, Text S1). AM does not satisfy either property.

### CLR transformation absorbs geometric averaging

Applying the CLR transformation, CLR(*x_i_*) = log(*x_i_*/GM(**x**)), to the same dataset (panel E of Figure **2**) produces per-sample, scale-invariant abundances centered on the WS-GM. Across-sample arithmetic means of CLR values are F1 = 0.10, F2 = −0.05, F3 = −0.06, ranking the three features in the same order as AS-GM on raw counts or proportions. This is not a coincidence: AS-AM on CLR-transformed values equals log(AS-GM) up to a group-specific additive constant, so CLR-based and GM-based rankings coincide exactly (formal statement in Text S1). CLR therefore provides the natural arithmetic space for compositional data: arithmetic averaging in CLR space is mathematically equivalent to geometric averaging in the original space, and differences of CLR values correspond to log fold-changes between features (CLR(*x_i_*) − CLR(*x_j_*) = log(*x_i_*/*x_j_*)), enabling interpretable cross-sample comparisons.

### Arithmetic operators are interpretable only on raw counts

On raw count data, the total sum has a clear biological meaning: it reflects sequencing depth (or extraction yield in metabolomics), and the AM is simply this total divided by the number of features. Once the data are normalized, however, the total sum becomes either a trivial constant (1 for proportions) or a quantity tied to the chosen reference feature (for ratios), and the AM loses any direct interpretation. The GM, by contrast, retains the same meaning across all representations: a central tendency of the multiplicative structure of the data, invariant to within-sample rescaling (Text S1).

### Empirical extent of AM/GM discordance in the gut microbiome

To establish whether the AM/GM discordance illustrated by the worked examples of generalizes to real data, we systematically computed AM- and GM-based rankings on relative proportions for all 102 prevalent genera in the dietswap baseline (Figure **3**). Within each dietary group, the overall agreement between AM- and GM-based rankings was high (Spearman *ρ* = 0.972 in AFR and 0.974 in AAM, panel A), yet this aggregate correlation conceals substantial local disagreement: 20 genera in AFR and 24 in AAM showed rank differences exceeding five positions between the two operators, with seven genera per group exceeding ten positions (panel B).

**FIG 3.**
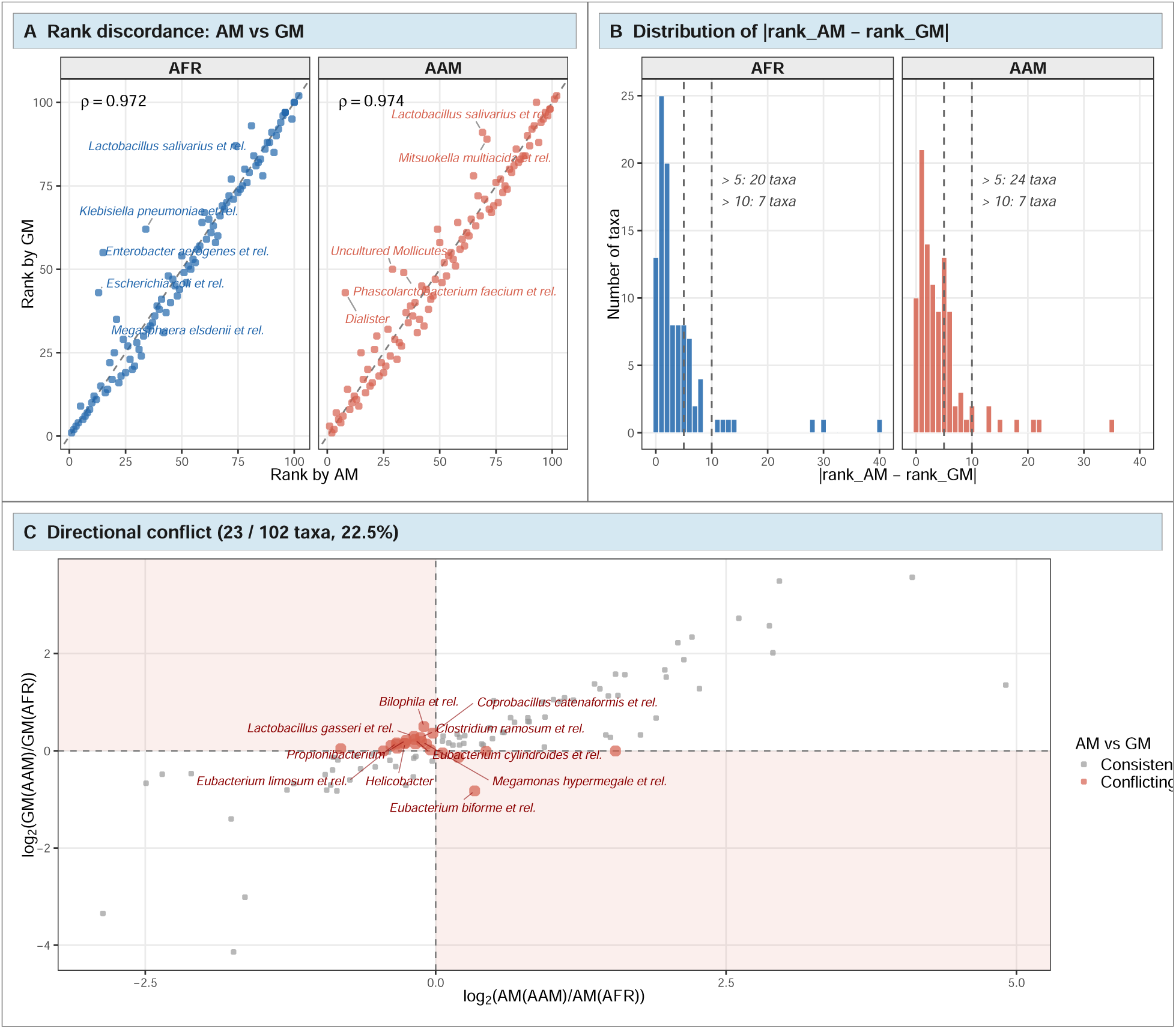
Empirical extent of AM/GM discordance across the gut microbiome on relative proportions. dietswap baseline, 102 prevalent genera; AM and GM computed across samples within each group on relative proportions, i.e. on counts normalized to the within-sample total. Panel A — scatter of GM-based rank (y-axis) versus AM-based rank (x-axis) for each genus, faceted by dietary group; the dashed diagonal corresponds to perfect concordance, and the inset reports the Spearman correlation between the two rankings. The five most discordant genera in each group are labelled. Panel B — distribution of |rank_AM − rank_GM| across genera, faceted by group; dashed vertical lines mark the discordance thresholds 5 and 10. Panel C — directional conflict between groups: each point is a genus plotted by log_2_(AM(AAM)/AM(AFR)) on the x-axis and log_2_(GM(AAM)/GM(AFR)) on the y-axis. The shaded second and fourth quadrants identify the 23 genera (22.5% of the panel) where AM and GM disagree on the direction of group-level enrichment; representative conflicting taxa are labelled.

More consequentially, 23 of the 102 genera (22.5%) exhibited directional conflict between groups: AM and GM disagreed on which dietary group displayed higher relative abundance (panel C, shaded quadrants). Conflicting taxa include genera of established biological relevance in diet–microbiome studies, such as *Bacteroides*, *Eubacterium* and *Bilophila*, whose group-level enrichment, and therefore their candidacy as dietary biomarkers, depends on the choice of averaging operator. The complete list of conflicting genera with operator-specific log_2_ fold-changes is provided in Table S2. This pattern is not an artefact of pseudocount handling: restricting the analysis to the 98 of 102 genera with non-zero counts in every baseline sample (where the pseudocount choice has no effect by construction) yielded a Spearman correlation *ρ* ≥ 0.9997 between AM- and GM-based rankings computed with and without pseudocount, and a directional-conflict percentage of 21.4% (with pseudocount) or 20.4% (without), within one percentage point of the full-panel result (Table S3).

In contrast, the Spearman correlation between GM-based and CLR-based rankings was numerically equal to 1.000 in both groups (Table S3), providing direct empirical confirmation of Proposition 2 (Text S1): AS-AM on CLR-space preserves the AS-GM ordering exactly. Together, these observations indicate that AM/GM discordance affects a substantial fraction of prevalent gut microbial genera: roughly one in five yields opposite group-level conclusions depending on whether arithmetic or geometric averaging is used to summarize relative abundances.

### Operator choice propagates from feature ranking to between-group differential inference

The analysis so far focused on feature ranking: we showed (Fig. 3) that AM- and GM-based summaries disagree on the direction of group-level enrichment for one in five prevalent genera. The same operator choice, however, also affects between-group differential inference, where log_2_ fold-changes and statistical tests are computed. To illustrate this propagation on real data, we selected three biologically relevant gut taxa from the same dietswap panel of 102 prevalent genera: F1 = *Bacteroides vulgatus et rel.*, F2 = *Clostridium cellulosi et rel.*, and F3 = *Sporobacter termitidis et rel.*. These taxa have mean proportions between 1.3% and 16% across groups (*n*_AFR_ = 17, *n*_AAM_ = 21), and were analyzed under the same four normalizations of Figure **1** (Figure **4** and Figure **5**). These three taxa were chosen automatically as the triple that maximizes ranking instability under AM across the four normalizations (selection procedure detailed in Methods); the goal is not to claim a specific biomarker, but to demonstrate concretely how the AM/GM divergence seen at the panel-wide level affects effect-size estimates and *p*-values for a representative case. The two dietary groups play the role of the two samples in the toy example: each table cell reports the AS-AM and AS-GM of per-sample values across all samples in the corresponding group.

**FIG 4.**
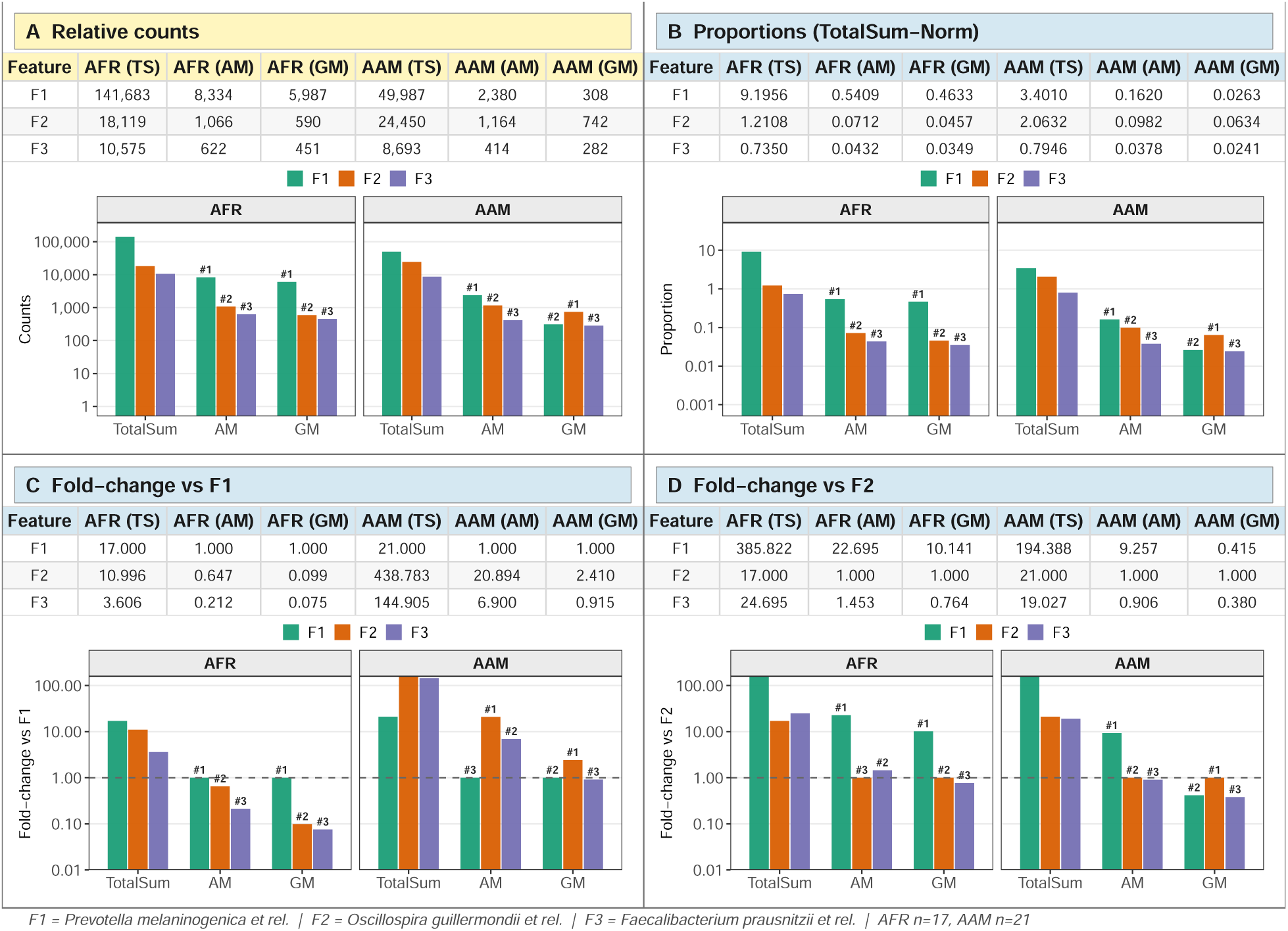
Empirical analog of Figure 1 on the dietswap dataset (baseline timepoint, n=38). Three taxa F1=*Bacteroides vulgatus et rel.*, F2=*Clostridium cellulosi et rel.*, F3=*Sporobacter termitidis et rel.* are summarized within each dietary group (AFR, AAM) under four within-sample normalizations. Panel A — relative counts. Panel B — proportions (TotalSum-Norm). Panel C — fold-change vs F1 (F1-Norm). Panel D — fold-change vs F2 (F2-Norm). Within each panel, the table reports the across-samples TotalSum, AM and GM per feature and group; the bar chart displays the same statistics on a log_10_ y-axis (linear-data, log-scale display). Bars are colored by feature using the Dark2 palette. Across panels, AM-based feature rankings change with the normalization choice (Proposition 1), whereas GM-based rankings remain invariant by the multiplicative property GM(*a*/*b*)=GM(*a*)/GM(*b*).

**FIG 5.**
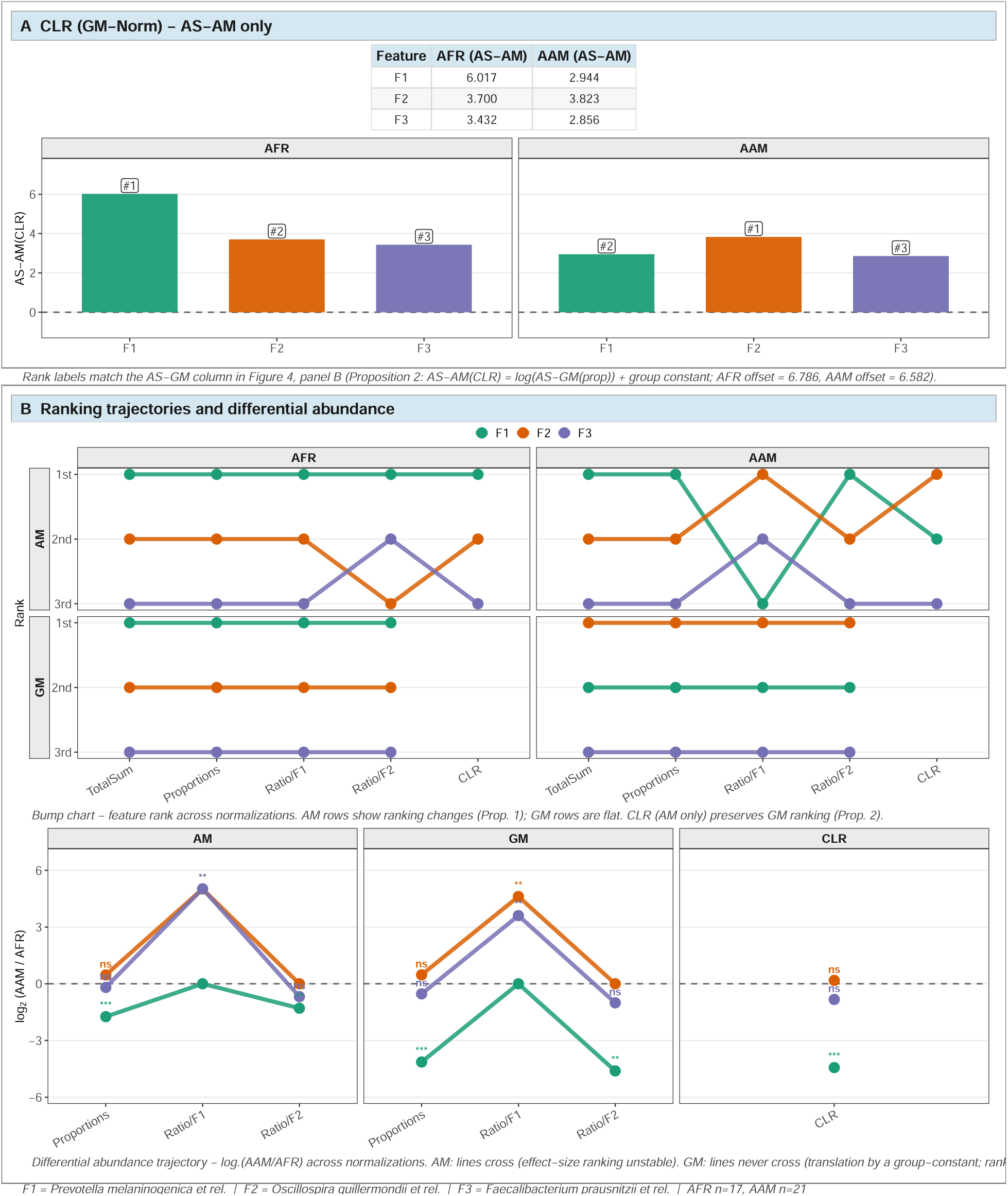
CLR representation, ranking trajectories, and operator-dependent differential inference for the three taxa of Figure 4. Panel A — AS-AM(CLR) per feature and group. Rank labels (#1, #2, #3) match the AS-GM column in Figure 4, panel B, since AS-AM(CLR)=log(AS-GM(prop))+*c_g_* (group offsets reported as a panel footnote). Panel B (top) — bump chart of feature rank across normalizations (TotalSum, Proportions, Ratio/F1, Ratio/F2, CLR), faceted by Statistic (rows: AM, GM) and Group (columns: AFR, AAM). AM rows show crossing lines, indicating ranking instability; GM rows show flat, non-crossing lines, illustrating Proposition 1 in visual form. The Spearman correlation between GM- and CLR-based rankings across all 102 prevalent genera equals 1.000 in both groups (Table S3), providing direct empirical confirmation of Proposition 2 of Text S1. Panel B (bottom) — log_2_ fold-change AAM/AFR across normalizations, faceted by operator (AM, GM, CLR). Under AM, log_2_FC values vary in unstructured ways and feature lines cross. Under GM, log_2_FC values vary by a group-constant translation and feature lines never cross, preserving the relative ordering of effects. CLR yields a single value per feature, anchored to the GM ranking. Asterisks denote Wilcoxon rank-sum significance (raw p-values: ∗ *p <* 0.05, ∗∗ *p <* 0.01, ∗ ∗ ∗ *p <* 0.001, ns = not significant). p-values depend on the data transformation (proportions, ratio, CLR) but not on the AM versus GM choice on the same data.

### Normalization dependence on real microbiome features

Already on raw counts (Figure **4**, panel A), AM and GM yield different rankings: AM is inflated by samples with extreme sequencing depth, whereas AS-GM provides a depth-robust estimate of typical abundance, mirroring the illustrative example of Figure **2**. The discrepancy widens after WS normalization: in panel B (proportions), AM ranks the three features differently between AFR and AAM, and the AM-based ranking changes again when proportions are replaced by fold-changes relative to F1 (panel C) or to F2 (panel D). The AS-GM column, by contrast, gives the same ordering of F1, F2, F3 across all four normalizations and in both groups, as guaranteed by Proposition 1 (Text S1). The practical consequence is that, given identical sequencing data, the AM-based identification of the most enriched taxon between AFR and AAM is reference-dependent, whereas the GM-based identification is not.

### CLR-space inference and operator-dependent fold-changes

The CLR-transformed values for the same three taxa (Figure **5**, panel A) confirm Proposition 2: within each group, AS-AM on CLR values preserves the AS-GM ranking of proportions, with group-specific offsets reported as a panel footnote. This is consistent with the exact identity *ρ*(rank_GM, rank_CLR) = 1.000 documented above for both groups across all 102 genera. CLR therefore absorbs the geometric-mean aggregation into the data transformation itself, leaving arithmetic averaging on CLR-space as the operator of choice.

The bump chart in panel B (top) summarizes the within-group ranking pattern: in the AM facets the lines connecting features across normalizations cross repeatedly, indicating ranking instability; in the GM facets the lines are flat and parallel, and the CLR column attached to the AM facets sits at the same rank values as the GM column. The log_2_-fold-change trajectory (panel B, bottom) extends the same logic to between-group comparison. Under AM, log_2_FC values vary in unstructured ways across normalizations and feature lines cross; under GM, log_2_FC values shift by a group-specific constant under reference change (a direct consequence of GM(*a*/*b*) = GM(*a*)/GM(*b*), Proposition 3 of Text S1), so feature lines never cross and the relative ordering of effects is preserved. CLR returns a single, reference-free value per feature, equivalent in ranking to GM. Wilcoxon rank-sum *p*-values, being rank-based, are identical for AM and GM under the same data transformation; they vary between proportions, ratios and CLR because the test is sensitive to the shape of the cross-sample distribution, not to the choice of summary operator.

These observations show that AM-based between-group inference is unstable in three respects simultaneously (effect magnitude, ranking of features by effect size, and statistical significance through the chosen transformation), whereas GM- and CLR-based inference is stable in the relevant sense. Consistent with the framing of the Introduction, this illustration is not a proposal for a new differential abundance test, but a demonstration that any downstream inferential procedure inherits the instability of its input summary; GM- or CLR-based summaries provide the only internally consistent foundation for such procedures.

## DISCUSSION

In gut microbiome and broader compositional NGS analyses, the choice of summary operator used to rank features within a group of samples is typically left implicit and defaults to the arithmetic mean. We have shown that this default has consequences.

Theoretically, AM-based rankings change with the within-sample normalization domain (Proposition 1, Text S1), so that the same dataset can yield different feature hierarchies depending on whether one summarizes raw counts, proportions, or ratios to a reference feature. Empirically, this instability translates into directional disagreement on real microbiome data: in the dietswap baseline, 23 of 102 prevalent genera (22.5%) yielded opposite group-level conclusions under AM and GM (Figure **3**), and the conflict involves taxa of established biological relevance for diet–microbiome interactions, including *Bacteroides*, *Eubacterium* and *Bilophila*. The geometric mean, by construction, is invariant to within-sample normalization (Proposition 1), satisfies an exact reciprocal identity under reference change (Proposition 3), and produces feature rankings that coincide exactly with those obtained by arithmetic averaging on CLR-transformed data (Proposition 2; Spearman *ρ* = 1.000 on all 102 genera in both groups).

These observations clarify the relationship between feature ranking and downstream differential abundance (DA) testing. Established DA methods such as ANCOM-BC [10] and LinDA [23] operate on geometric-mean or log-ratio reference structures and address the inferential question of whether feature abundances differ significantly between groups. The problem analyzed here is logically prior: ensuring that the aggregation operator used to summarize feature abundances within a group is itself consistent with the compositional nature of NGS data. AM-based summaries violate this consistency, and any inferential procedure that ingests an AM-based feature ranking inherits its instability. The three-taxa illustration in Figure **5** makes this propagation explicit: under AM, log_2_ fold-changes vary with the chosen reference and the relative ordering of features by effect size is not preserved; under GM, fold-changes shift by a group-constant translation under reference change and their relative ordering is preserved; under CLR, a single reference-free value per feature returns the same ranking as GM.

The practical implication for microbiome and genomics research is that biomarker rankings, signatures, and effect-size comparisons should be derived from GM-based summaries on raw counts or proportions, or equivalently from arithmetic averaging on CLR-transformed data. This recommendation requires no additional modeling assumptions and no new computational tools, and is fully compatible with existing DA pipelines, which already adopt log-ratio or geometric-mean reference frames internally. By removing operator-induced instability at the summarization stage, GM- and CLR-based pipelines provide a reproducible foundation for cross-study comparisons and meta-analyses, including the kind of microbial signature analyses we have previously reported in clinical microbiome contexts [16, 17, 18].

The framework developed here applies to the strictly positive subset of compositional data, the regime where AM and GM aggregation differ. NGS datasets typically contain structural and sampling zeros, and zero-handling strategies (pseudocounts, zero-inflated models, non-parametric estimators) remain orthogonal to the operator-choice problem analyzed here. The arithmetic sum on raw counts retains its biological meaning as an estimate of total sequencing signal allocated to a feature, but loses this interpretation once the data are normalized (Section S1.5 of Text S1). Future work integrating the GM/CLR framework with sparsity-aware models would extend its applicability without altering its core compositional logic.

## MATERIALS AND METHODS

### Worked-example datasets

The illustrative datasets used in Figure **1** and Figure **2** are synthetic compositional matrices adapted from the worked examples in **(author?)** [15], constructed to expose the compositional properties analyzed in this work, and do not represent any real biological measurement. Figure **1** uses three features (F1–F3) measured in two samples (S1, S2), with integer counts chosen so that AM-based rankings of feature ratios change across normalizations whereas AS-GM rankings remain invariant. Figure **2** extends the construction to five samples with sequencing depths spanning three orders of magnitude (S2 = 223 reads, S4 = 138,976 reads), demonstrating the same properties under extreme depth heterogeneity. Panel E of Figure **2** additionally applies the centered log-ratio transformation to that same dataset.

### Empirical dataset and preprocessing

To validate the theoretical framework on real compositional data, we analyzed the publicly available dietswap dataset [24], accessible via the microbiome R package (v. 1.32.0). The dataset comprises 16S rRNA amplicon sequencing data from 222 fecal samples collected from rural African (AFR) and African-American (AAM) individuals undergoing a two-week dietary exchange. Analysis was restricted to the baseline timepoint (n = 38 samples; AFR n = 17, AAM n = 21) to avoid within-subject correlation across timepoints. Taxa with prevalence below 50% in at least one group were excluded, retaining 102 genera for compositional analysis. A unit pseudocount (+1) was added to all retained counts to ensure strict positivity prior to log-based transformations. The choice of pseudocount = 1 is principled rather than arbitrary on count-based NGS data: it equals the smallest count observable on a discrete sequencing platform, and avoids introducing any a priori assumption about the relative abundance of features absent in a given sample, an assumption that in practice cannot be informed from the data itself. Pseudocount values smaller or larger than 1 implicitly carry such priors and are therefore not pursued. To verify that the AM/GM discordance pattern is not an artefact of pseudocount handling, all ranking and conflict statistics were recomputed on the subset of 98 genera with non-zero counts in every baseline sample (where pseudocount handling is mathematically irrelevant); the results, reported in Table S3, agree with the full-panel analysis to within one percentage point. Relative proportions were obtained by sample-wise normalization to the total counts.

### Compositional aggregation and CLR transformation

For each feature *i* across *n* samples within a group, the across-sample arithmetic and geometric means were computed as AS-AM(**x***_i_*) = (1/*n*) ∑*_k_ x_i_*_,*k*_ and AS-GM(**x***_i_*) = (∏*_k_ x_i_*_,*k*_)^1/*n*^, respectively. Within-sample normalizations applied to the empirical and worked-example datasets included: (i) raw counts; (ii) proportions, obtained by dividing each count by its within-sample total; and (iii) ratios to a chosen reference feature (F1 or F2 in the worked examples; the corresponding genera in the three-taxa empirical illustration). The centered log-ratio transformation was computed sample-wise as CLR(*x_i_*_,*k*_) = log(*x_i_*_,*k*_/WS-GM(**x**_·,*k*_)), where WS-GM(**x**_·,*k*_) is the geometric mean over all features in sample *k*. Formal statements and proof sketches of the underlying invariance properties are provided in Text S1.

### Empirical validation procedures

The empirical validation comprised two complementary analyses. First, a systematic comparison of AM- and GM-based rankings was performed on all 102 prevalent genera within each dietary group, computing (i) Spearman correlation between AM and GM rankings on relative proportions; (ii) the distribution of |rank_AM − rank_GM| across genera; (iii) cross-group directional conflict, defined as the disagreement between AM and GM on which group exhibits higher relative abundance. The same pipeline was repeated for GM-based versus CLR-based rankings. The complete list of conflicting genera is provided in Table S2.

Second, three taxa were selected from the pool of features with mean proportion ≥ 1% in both dietary groups, choosing the triple that maximized the number of distinct AM-based feature rankings across the four within-sample normalizations (raw counts, proportions, ratio to F1, ratio to F2) within each group. This selection criterion identifies the most illustrative case of normalization dependence under AM; it does not affect GM-based rankings, which are guaranteed invariant by Proposition 1 (Text S1) and were verified numerically. For the differential illustration in Figure **5**, log_2_ fold-changes (AAM/AFR) were computed under each operator × normalization combination, and Wilcoxon rank-sum tests were performed on the corresponding per-sample values.

### Software and reproducibility

All analyses were performed in R (v. 4.5.0) using the phyloseq, microbiome, dplyr and ggplot2 packages. The R scripts used to generate all Figures are provided as File S1.

## ACKNOWLEDGMENTS

The authors thank the developers of the microbiome R package and the authors of the dietswap dataset for making the data publicly available.

## DATA AVAILABILITY STATEMENT

This study analyzed the publicly available dietswap dataset [24], accessible through the microbiome R package (v. 1.32.0; Bioconductor). No new sequencing data were generated. R analysis scripts and the filtered proportion matrix are provided as File S1. Tables S1, S2 and S3 are provided as supplementary materials accompanying this article.

## DECLARATION OF GENERATIVE AI USE IN THE WRITING PROCESS

During the preparation of this work, the authors used ChatGPT (OpenAI) and Claude (Anthropic) for language editing, grammar correction, and improvement of readability. After using these tools, the authors reviewed and edited the content as needed and take full responsibility for the content of the publication. No AI tools were used to generate or manipulate research data, results, or figures.

## References

[1] Gloor, G. ALDEx2: ANOVA-Like Differential Expression tool for compositional data. ALDEX Manual Modular, 20, 1–11 (2015).

[2] Gloor, G. B., Macklaim, J. M., Pawlowsky-Glahn, V., & Egozcue, J. J. Microbiome datasets are compositional: and this is not optional. Frontiers in Microbiology, 8, 2224 (2017).

[3] Patrick D Schloss. Rarefaction is currently the best approach to control for uneven sequencing effort in amplicon sequence analyses. Msphere, pages e00354–23, 2024.

[4] Patrick D Schloss. Waste not, want not: revisiting the analysis that called into question the practice of rarefaction. Msphere, 9(1):e00355–23, 2024.

[5] Michael Greenacre, Eric Grunsky, John Bacon-Shone, Ionas Erb, and Thomas Quinn. Aitchison’s compositional data analysis 40 years on: A reappraisal. Statistical Science, 38(3):386–410, 2023.

[6] John Aitchison. The statistical analysis of compositional data. Journal of the Royal Statistical Society: Series B (Methodological), 44(2):139–160, 1982.

[7] Beibei Wang, Fengzhu Sun, and Yihui Luan. Comparison of the effectiveness of different normalization methods for metagenomic cross-study phenotype prediction under heterogeneity. Scientific Reports, 14(1):7024, 2024.

[8] M Luz Calle, Meritxell Pujolassos, and Antoni Susin. coda4microbiome: compositional data analysis for microbiome cross-sectional and longitudinal studies. BMC bioinformatics, 24(1):82, 2023.

[9] Huang Lin and Shyamal Das Peddada. Multigroup analysis of compositions of microbiomes with covariate adjustments and repeated measures. Nature Methods, 21(1):83–91, 2024.

[10] Huang Lin and Shyamal Das Peddada. Analysis of compositions of microbiomes with bias correction. Nature communications, 11(1):3514, 2020.

[11] Huang Lin and Shyamal Das Peddada. Analysis of microbial compositions: a review of normalization and differential abundance analysis. NPJ biofilms and microbiomes, 6(1):1–13, 2020.

[12] Dionne Swift, Kellen Cresswell, Robert Johnson, Spiro Stilianoudakis, and Xingtao Wei. A review of normalization and differential abundance methods for microbiome counts data. Wiley Interdisciplinary Reviews: Computational Statistics, 15(1):e1586, 2023.

[13] James T Morton, Clarisse Marotz, Alex Washburne, Justin Silverman, Livia S Zaramela, Anna Edlund, Karsten Zengler, and Rob Knight. Establishing microbial composition measurement standards with reference frames. Nature communications, 10(1):1–11, 2019.

[14] Yingtian Hu, Glen A Satten, and Yi-Juan Hu. Locom: A logistic regression model for testing differential abundance in compositional microbiome data with false discovery rate control. Proceedings of the National Academy of Sciences, 119(30):e2122788119, 2022.

[15] Philip J Fleming and John J Wallace. How not to lie with statistics: the correct way to summarize benchmark results. Communications of the ACM, 29(3):218–221, 1986.

[16] Giorgia Renga, Emilia Nunzi, Claudia Stincardini, Marilena Pariano, Matteo Puccetti, Giuseppe Pieraccini, Claudia Di Serio, Maurizio Fraziano, Noemi Poerio, Vasileios Oikonomou, et al. Cpx-351 exploits the gut microbiota to promote mucosal barrier function, colonization resistance, and immune homeostasis. Blood, 143(16):1628–1645, 2024.

[17] Claudio Costantini, Emilia Nunzi, Angelica Spolzino, Melissa Palmieri, Giorgia Renga, Teresa Zelante, Lukas Englmaier, Katerina Coufalikova, Zdeněk Spáčil, Monica Borghi, et al. Pharyngeal microbial signatures are predictive of the risk of fungal pneumonia in hematologic patients. Infection and immunity, 89(8), 2021.

[18] Giorgia Renga, Fiorella D’Onofrio, Marilena Pariano, Roberta Galarini, Carolina Barola, Claudia Stincardini, Marina M Bellet, Helmut Ellemunter, Cornelia Lass-Flörl, Claudio Costantini, et al. Bridging of host-microbiota tryptophan partitioning by the serotonin pathway in fungal pneumonia. Nature Communications, 14(1):5753, 2023.

[19] M Luz Calle. Statistical analysis of metagenomics data. Genomics & informatics, 17(1), 2019.

[20] M Luz Calle and Antoni Susin. coda4microbiome: compositional data analysis for microbiome studies. bioRxiv, 2022.

[21] Thomas P Quinn, Ionas Erb, Mark F Richardson, and Tamsyn M Crowley. Understanding sequencing data as compositions: an outlook and review. Bioinformatics, 34(16):2870–2878, 2018.

[22] Paul J McMurdie and Susan Holmes. Waste not, want not: why rarefying microbiome data is inadmissible. PLoS Computational Biology, 10(4):e1003531, 2014.

[23] Huijuan Zhou, Kejun He, Jun Chen, and Xianyang Zhang. LinDA: linear models for differential abundance analysis of microbiome compositional data. Genome Biology, 23(1):95, 2022.

[24] S. J. D. O’Keefe, J. V. Li, L. Lahti, J. Ou, F. Carbonero, K. Mohammed, J. M. Posma, J. Kinross, E. Wahl, E. Ruder, et al., Fat, fibre and cancer risk in African Americans and rural African. Nature Communications, vol. 6, no. 1, p. 6342, 2015.

