## Supplementary file for "Geometric averaging provides normalization-invariant feature ranking in compositional sequencing data"

E. Nunzi, L. Romani

#### Contents

**Text S1 — Mathematical framework** **2**

**Table S1 — Pairwise ratio comparisons (Fig. 2 worked example)** **4**

**Table S2 — Directional conflict between AM and GM** **4**

**Table S3 — Sensitivity to pseudocount choice** **6**

**File S1 — R analysis scripts** **7**

**Contents.** This document collects the supplementary materials cited in the main text:

- **Text S1** — Formal statements and proof sketches of the three propositions on AM/GM aggregation under within-sample normalization.
- **Table S1** — Pairwise ratio comparisons on the five-sample worked-example dataset of Figure 2, illustrating the failure of the reciprocal property under AM and its exact validity under GM.
- **Table S2** — Genera with directional conflict between AM- and GM-based group enrichment (full list, n=23). Provided as a separate CSV file (`Table_S2_directional_conflict.csv`) generated by the R analysis script `fig3.R`.
- **Table S3** — Sensitivity of the AM/GM discordance pattern to pseudocount handling (zero-free subset analysis), and concordance between GM- and CLR-based rankings.
- **File S1** — R analysis scripts (`fig1.R` . . . `fig5.R`, `sensitivity_pseudocount.R`), provided as separate files.

### Text S1 — Mathematical framework

This Supporting Text provides the formal statements and proof sketches underlying the analyses in the Results section of the main text (subsection "Geometric mean provides normalization-invariant feature ranking in compositional data").

#### S1.1 Setup and notation

Let $\mathbf{X} = (x_{i,k})$ be a $D \times n$ matrix of strictly positive abundances, where $i = 1, \dots, D$ indexes features (e.g., genera, genes) and $k = 1, \dots, n$ indexes samples within a group. We denote by $\mathbf{x}_i = (x_{i,1}, \dots, x_{i,n})$ the vector of abundances of feature $i$ across samples, and by $\mathbf{x}_{\cdot,k} = (x_{1,k}, \dots, x_{D,k})$ the vector of abundances in sample $k$.

The aggregation operators used throughout the paper are:

$$\text{AS-AM}(\mathbf{x}_i) = \frac{1}{n}\sum_{k=1}^{n} x_{i,k}, \qquad \text{AS-GM}(\mathbf{x}_i) = \left(\prod_{k=1}^{n} x_{i,k}\right)^{1/n} = \exp\left(\frac{1}{n}\sum_{k=1}^{n}\log x_{i,k}\right),$$

$$\text{WS-GM}(\mathbf{x}_{\cdot,k}) = \left(\prod_{i=1}^{D} x_{i,k}\right)^{1/D}.$$

The CLR transformation is defined sample-wise as

$$\text{CLR}(x_{i,k}) = \log\frac{x_{i,k}}{\text{WS-GM}(\mathbf{x}_{\cdot,k})} = \log x_{i,k} - \frac{1}{D}\sum_{j=1}^{D}\log x_{j,k}.$$

A within-sample (WS) normalization is any transformation $y_{i,k} = x_{i,k}/N_k$ where $N_k > 0$ depends on the sample $k$ but not on the feature $i$. Special cases include $N_k = \sum_j x_{j,k}$ (proportions) and $N_k = x_{j,k}$ for some fixed reference feature $j$ (ratios to a reference).

#### S1.2 Proposition 1: Ranking invariance under within-sample normalization

**Statement.** For any choice of WS normalization factors $\{N_k\}_{k=1}^{n}$ and any pair of features $i, j$:

$$\frac{\text{AS-GM}(\mathbf{y}_i)}{\text{AS-GM}(\mathbf{y}_j)} = \frac{\text{AS-GM}(\mathbf{x}_i)}{\text{AS-GM}(\mathbf{x}_j)}. \tag{1}$$

In particular, the feature ranking induced by AS-GM is identical on $\mathbf{X}$ and on the WS-normalized matrix $\mathbf{Y}$. The analogous identity does *not* hold for AS-AM in general: when $N_k$ varies across samples, the ratio $\text{AS-AM}(\mathbf{y}_i)/\text{AS-AM}(\mathbf{y}_j)$ depends on the joint distribution of $x_{i,k}$, $x_{j,k}$ and $N_k$, and the AS-AM-induced ranking can differ between $\mathbf{X}$ and $\mathbf{Y}$.

**Proof sketch.** For AS-GM,

$$\text{AS-GM}(\mathbf{y}_i) = \left(\prod_{k=1}^{n}\frac{x_{i,k}}{N_k}\right)^{1/n} = \frac{(\prod_k x_{i,k})^{1/n}}{(\prod_k N_k)^{1/n}} = \frac{\text{AS-GM}(\mathbf{x}_i)}{G},$$

where $G = (\prod_k N_k)^{1/n}$ is a feature-independent constant. Taking the ratio over two features cancels $G$, yielding (1).

For AS-AM, $\text{AS-AM}(\mathbf{y}_i) = (1/n)\sum_k x_{i,k}/N_k$ is a weighted average of $x_{i,k}$ with weights $1/N_k$ that are feature-independent but sample-dependent. The ratio $\text{AS-AM}(\mathbf{y}_i)/\text{AS-AM}(\mathbf{y}_j)$ therefore depends on how mass is distributed across samples, and a finite counterexample suffices to show that ranking is not preserved. Such a counterexample is provided in Figure 1 of the main text (panel D vs. panels A–C). □

#### S1.3 Proposition 2: CLR–AM equivalence to log of AS-GM

**Statement.** For each feature $i$,

$$\text{AS-AM}(\text{CLR}(\mathbf{x}_i)) = \log\bigl(\text{AS-GM}(\mathbf{x}_i)\bigr) - c, \tag{2}$$

where $c = (1/n)\sum_{k=1}^{n} \log \text{WS-GM}(\mathbf{x}_{\cdot,k})$ is a constant that depends on the group of samples but not on the feature $i$. Consequently, the feature ranking induced by AS-AM on CLR-transformed data coincides exactly with the ranking induced by AS-GM on the original data.

**Proof.** By definition,

$$\begin{aligned}
\text{AS-AM}(\text{CLR}(\mathbf{x}_i)) &= \frac{1}{n}\sum_{k=1}^{n} [\log x_{i,k} - \log \text{WS-GM}(\mathbf{x}_{\cdot,k})] \\
&= \frac{1}{n}\sum_{k=1}^{n} \log x_{i,k} - \frac{1}{n}\sum_{k=1}^{n} \log \text{WS-GM}(\mathbf{x}_{\cdot,k}) \\
&= \log\bigl(\text{AS-GM}(\mathbf{x}_i)\bigr) - c,
\end{aligned}$$

which is (2). Subtracting the same identity for two features $i, j$ removes $c$:

$$\text{AS-AM}(\text{CLR}(\mathbf{x}_i)) - \text{AS-AM}(\text{CLR}(\mathbf{x}_j)) = \log\left(\frac{\text{AS-GM}(\mathbf{x}_i)}{\text{AS-GM}(\mathbf{x}_j)}\right).$$

Because the logarithm is strictly monotonic, the ordering of features by AS-AM(CLR) coincides with the ordering by AS-GM. □

#### S1.4 Proposition 3: Exact reciprocal property of GM under reference change

**Statement.** For any two features $i, j$,

$$\text{AS-GM}\left(\frac{\mathbf{x}_i}{\mathbf{x}_j}\right) = \frac{\text{AS-GM}(\mathbf{x}_i)}{\text{AS-GM}(\mathbf{x}_j)}, \qquad \text{AS-GM}\left(\frac{\mathbf{x}_j}{\mathbf{x}_i}\right) = \frac{1}{\text{AS-GM}(\mathbf{x}_i/\mathbf{x}_j)}.$$

The corresponding identities do not hold for AS-AM: in general,

$$\text{AS-AM}(\mathbf{x}_i/\mathbf{x}_j) \neq \frac{\text{AS-AM}(\mathbf{x}_i)}{\text{AS-AM}(\mathbf{x}_j)}, \qquad \text{AS-AM}(\mathbf{x}_i/\mathbf{x}_j) \neq \frac{1}{\text{AS-AM}(\mathbf{x}_j/\mathbf{x}_i)}.$$

**Proof.** Direct application of the multiplicative property:

$$\left(\prod_k x_{i,k}/x_{j,k}\right)^{1/n} = \left(\prod_k x_{i,k}\right)^{1/n} \Big/ \left(\prod_k x_{j,k}\right)^{1/n}.$$

The reciprocal identity follows by substitution. The failure of the analogous AM identities is empirically illustrated in Table S1 below (rows F1/F2 and F2/F1: AM ratios $1.32 \neq 1/1.02 = 0.98$). □

#### S1.5 Remark: Arithmetic sums are interpretable only on raw counts

The arithmetic sum $\sum_k x_{i,k}$ over raw counts represents the total sequencing signal allocated to feature $i$ across all samples, a quantity proportional to total read depth and, under appropriate experimental controls (spike-ins, equimolar libraries), to total feature-specific signal. After WS normalization, the analogous sum $\sum_k y_{i,k} = \sum_k x_{i,k}/N_k$ no longer corresponds to a total signal, because each term lives in a different scale (one per sample) determined by $N_k$. The sum therefore loses its biological interpretation. The arithmetic mean $\text{AS-AM}(\mathbf{y}_i) = (1/n)\sum_k y_{i,k}$ inherits this loss of interpretation and, as shown in Proposition 1, fails to preserve feature ranking across normalizations. By contrast, AS-GM remains interpretable on any positive normalization of the data, because Proposition 1 guarantees that the ranking it induces is the same as on raw counts.

#### Table S1 — Pairwise ratio comparisons on the worked-example dataset of Figure 2

Across-sample arithmetic and geometric means of the six pairwise ratios $F_i/F_j$ on the five-sample worked-example dataset of Figure 2 of the main text. The two right-most columns illustrate the reciprocal property under each operator: the geometric mean satisfies $\mathrm{GM}(F_i/F_j)\cdot\mathrm{GM}(F_j/F_i) = 1$ exactly (Proposition 3), whereas the corresponding product for the arithmetic mean exceeds 1 systematically by Cauchy–Schwarz, demonstrating that AM-based fold-changes are not internally consistent under reference change.

| **Pairwise ratio** | **AS-AM** | **AS-GM** | **AM**$(F_i/F_j)\cdot$ **AM**$(F_j/F_i)$ | **GM**$(F_i/F_j)\cdot$ **GM**$(F_j/F_i)$ |
|---|---|---|---|---|
| $F_1/F_2$ | 1.320 | 1.166 | 1.339 | 1.000 |
| $F_2/F_1$ | 1.015 | 0.857 | | |
| $F_1/F_3$ | 1.495 | 1.177 | 1.603 | 1.000 |
| $F_3/F_1$ | 1.072 | 0.850 | | |
| $F_2/F_3$ | 1.092 | 1.009 | 1.180 | 1.000 |
| $F_3/F_2$ | 1.081 | 0.991 | | |

**Remark.** The exact equality $\mathrm{GM}(F_i/F_j)\cdot\mathrm{GM}(F_j/F_i) = 1$ is a direct consequence of $\mathrm{GM}(a/b) = \mathrm{GM}(a)/\mathrm{GM}(b)$ (Proposition 3, Text S1). The deviation of the AM-based product from 1 quantifies the amount by which AM-based fold-change estimates between two features fail to be mutually consistent under reference change.

ewpage

#### Table S2 — Genera with directional conflict between AM and GM

The complete list of the 23 prevalent genera (out of 102) for which AM- and GM-based group summaries disagree on the direction of relative abundance between AFR and AAM dietary groups, sorted by decreasing $|\log_2 \mathrm{FC_{GM}}|$. For each genus the table reports the AM and GM of relative proportions in each group (in units of $10^{-3}$), the direction of group-level enrichment under each operator (by construction always opposite), and the $\log_2$ fold-change AAM/AFR under each operator (the sign mismatch between the last two columns is the formal definition of directional conflict). The same data are provided as a separate CSV file (`Table_S2_directional_conflict.csv`) generated by the R analysis script `fig4.R`.

| **Genus** | **Mean proportion** ($\times 10^{-3}$) | | | | **Higher in** | | $\mathbf{log_2}$**FC AAM/AFR** | |
|---|---|---|---|---|---|---|---|---|
| | $\mathbf{AM_{AFR}}$ | $\mathbf{AM_{AAM}}$ | $\mathbf{GM_{AFR}}$ | $\mathbf{GM_{AAM}}$ | **AM** | **GM** | **AM** | **GM** |
| *Eubacterium biforme et rel.* | 2.16 | 2.73 | 1.37 | 0.78 | AAM | AFR | +0.336 | -0.824 |
| *Bilophila et rel.* | 0.26 | 0.24 | 0.16 | 0.23 | AFR | AAM | -0.102 | +0.501 |
| *Coprobacillus catenaformis et rel.* | 0.41 | 0.40 | 0.28 | 0.36 | AFR | AAM | -0.024 | +0.361 |
| *Lactobacillus gasseri et rel.* | 1.03 | 0.91 | 0.65 | 0.80 | AFR | AAM | -0.187 | +0.297 |
| *Clostridium ramosum et rel.* | 0.34 | 0.31 | 0.24 | 0.30 | AFR | AAM | -0.131 | +0.273 |
| *Eubacterium cylindroides et rel.* | 0.33 | 0.29 | 0.24 | 0.28 | AFR | AAM | -0.180 | +0.232 |
| *Propionibacterium* | 0.23 | 0.19 | 0.16 | 0.18 | AFR | AAM | -0.254 | +0.219 |

| **Genus** | **Mean proportion ($\times 10^{-3}$)** | | | | **Higher in** | | **$\log_2$FC AAM/AFR** | |
|---|---|---|---|---|---|---|---|---|
| | **AM$_{AFR}$** | **AM$_{AAM}$** | **GM$_{AFR}$** | **GM$_{AAM}$** | **AM** | **GM** | **AM** | **GM** |
| *Megamonas hypermegale et rel.* | 0.20 | 0.18 | 0.15 | 0.18 | AFR | AAM | -0.121 | +0.190 |
| *Eubacterium limosum et rel.* | 0.23 | 0.18 | 0.16 | 0.18 | AFR | AAM | -0.336 | +0.164 |
| *Helicobacter* | 0.33 | 0.28 | 0.24 | 0.27 | AFR | AAM | -0.265 | +0.153 |
| *Peptostreptococcus micros et rel.* | 0.34 | 0.28 | 0.24 | 0.27 | AFR | AAM | -0.268 | +0.153 |
| *Corynebacterium* | 0.20 | 0.18 | 0.15 | 0.17 | AFR | AAM | -0.174 | +0.142 |
| *Oceanospirillum* | 0.28 | 0.26 | 0.20 | 0.22 | AFR | AAM | -0.079 | +0.139 |
| *Campylobacter* | 0.56 | 0.45 | 0.39 | 0.43 | AFR | AAM | -0.337 | +0.137 |
| *Papillibacter cinnamivorans et rel.* | 2.39 | 2.73 | 1.79 | 1.63 | AAM | AFR | +0.196 | -0.135 |
| *Bacillus* | 0.23 | 0.18 | 0.16 | 0.17 | AFR | AAM | -0.385 | +0.108 |
| *Vibrio* | 0.37 | 0.30 | 0.27 | 0.28 | AFR | AAM | -0.332 | +0.052 |
| *Streptococcus bovis et rel.* | 4.21 | 4.39 | 2.49 | 2.41 | AAM | AFR | +0.059 | -0.045 |
| *Uncultured Bacteroidetes* | 0.44 | 0.25 | 0.20 | 0.21 | AFR | AAM | -0.817 | +0.042 |
| *Eubacterium siraeum et rel.* | 0.36 | 0.35 | 0.29 | 0.30 | AFR | AAM | -0.044 | +0.018 |
| *Streptococcus mitis et rel.* | 2.57 | 3.48 | 1.78 | 1.77 | AAM | AFR | +0.434 | -0.013 |
| *Uncultured Mollicutes* | 2.08 | 6.08 | 1.49 | 1.49 | AAM | AFR | +1.548 | -0.002 |
| *Fusobacteria* | 0.68 | 0.50 | 0.48 | 0.48 | AFR | AAM | -0.448 | +0.000 |

Genera are sorted by decreasing $|\log_2 \mathrm{FC}_{\mathrm{GM}}|$ (largest GM-based effect first). Columns "Higher in" report the dietary group enriched under each operator; by construction the AM and GM columns disagree for every row (this is the definition of directional conflict). Note that several conflicts involve genera with very small GM-based fold-change but a large AM-based fold-change driven by high-depth samples (e.g. Uncultured Mollicutes: $\log_2 \mathrm{FC}_{\mathrm{AM}} = +1.548$, $\log_2 \mathrm{FC}_{\mathrm{GM}} = -0.002$), illustrating how AM-based biomarker selection can be misled by depth-driven outliers.

### Table S3 — Sensitivity to pseudocount choice

Robustness of the AM/GM discordance pattern to pseudocount handling. The main analysis was performed on the full panel of 102 prevalent genera with a unit pseudocount (the smallest count observable on a discrete sequencing platform; see Materials and Methods for the rationale). To rule out artefacts induced by the pseudocount, we recomputed all ranking and conflict statistics on the subset of 98 genera with non-zero counts in every baseline sample, where pseudocount handling is mathematically irrelevant. The two analyses agree to within one percentage point in the directional-conflict count, and the Spearman correlation between AM- and GM-based rankings computed with and without pseudocount on the zero-free subset is essentially 1.

| **Analysis** | $N_g$ | $N_{\mathbf{conf}}$ | **% conf** | **Spearman** $\rho$ |
|---|---|---|---|---|
| Full panel, pseudo = 1 *(main)* | 102 | 23 | 22.5% | — |
| Zero-free, pseudo = 1 | 98 | 21 | 21.4% | — |
| Zero-free, no pseudo | 98 | 20 | 20.4% | — |
| *Concordance on zero-free subset (pseudo = 1 vs no pseudo):* | | | | |
| AM ranking | 98 | — | — | 0.9999 |
| GM ranking | 98 | — | — | 0.9997 |
| *GM- vs CLR-based ranking (full panel, pseudo = 1):* | | | | |
| AFR group | 102 | — | — | 1.0000 |
| AAM group | 102 | — | — | 1.0000 |

**Direction-conflict label agreement** between pseudocount = 1 and no-pseudocount analyses on the zero-free subset: 97/98 = 99.0%.

**Interpretation.** Across the three pseudocount analyses, the percentage of directional-conflict genera lies in the narrow range 20.4–22.5% (full range 2.1 percentage points), and the Spearman correlation between ranks computed with and without pseudocount on the zero-free subset exceeds 0.999 for both AM and GM. The AM/GM discordance reported in the main text is therefore a property of the data, not an artefact of zero handling. The exact equality (to four decimal places) of GM- and CLR-based rankings in both dietary groups provides empirical confirmation of Proposition 2 (Text S1): across-sample arithmetic means of CLR values are mathematically equivalent (up to an additive group-specific constant) to logarithms of across-sample geometric means of proportions, and therefore induce identical feature orderings.

ewpage

#### File S1 — R analysis scripts

The R scripts used to generate all figures and supplementary tables of the main text are provided as separate files alongside this PDF. Each script is self-contained: it loads the dietswap dataset (or reconstructs the worked-example data of Figures 1–2), performs the analysis, and writes the corresponding output(s).

**Worked-example illustrations:**

- `fig1.R` — two-sample illustration (Figure 1). Output: `F1.png`, `F1.pdf`.
- `fig2.R` — five-sample worked-example matrix with depth heterogeneity, including CLR panel E (Figure 2). Output: `F2.png`, `F2.pdf`.

**Empirical analyses on the dietswap dataset:**

- `fig3.R` — pervasiveness analysis on 102 prevalent genera (Figure 3). Output: `F3.png`, `F3.pdf`, and `Table_S2_directional_conflict.csv`.
- `fig4.R` — empirical analog of Figure 1 on three selected taxa (Figure 4). Output: `F4.png`, `F4.pdf`.
- `fig5.R` — CLR + ranking trajectories + differential inference (Figure 5). Output: `F5.png`, `F5.pdf`.

**Sensitivity analysis:**

- `sensitivity_pseudocount.R` — robustness of the AM/GM discordance to pseudocount handling (Table S3). Output: `Table_S3_pseudocount_sensitivity.csv`.

**Software environment.** All analyses were performed in R (v. 4.5.0) with the following CRAN and Bioconductor packages: `phyloseq`, `microbiome`, `dplyr`, `tidyr`, `ggplot2`, `scales`, `gridExtra`, `grid`, `gtable`, `RColorBrewer` and `ggrepel`.
